# Asymmetric gene flow maintains range edges in a marine invertebrate

**DOI:** 10.1101/2024.11.04.621918

**Authors:** Zachary B. Hancock, Nicole E. Adams, Perry L. Wood, Gideon S. Bradburd

## Abstract

Species are restricted to finite ranges, and when they are in demographic equilibrium with the environment this implies a failure to adapt at the range periphery. There are several competing evolutionary hypotheses for the maintenance and persistence of range edges. One hypothesis is gene swamping of edge populations, in which migrants from the center carrying alleles adapted to the range center but maladaptive along the edge hamper local adaptation. A competing explanation is dispersal limitation, in which reduced migration to the periphery keeps edge populations small, causing drift to dominate over selection. It has proven difficult to assess these potential generative scenarios due to the inherent complexity of species ranges. In this study, we leverage range-wide genomic sequencing data to investigate the evolution of range edges in an empirical system - *Haustorius canadensis*, the Canadian shovel bug. We find increased genetic drift and inbreeding on the range edges and detect signatures of asymmetric migration towards the range center. The inferred directionality of migration is consistent with the direction of the dominant current regimes in the North Atlantic. We conclude that anisotropic dispersal towards the range center leaves progressively fewer migrants to replenish peripheral populations, consistent with the migration limitation hypothesis, and generates the appearance of an abundant center.

## 1 INTRODUCTION

All species are restricted to finite ranges. How range edges form and persist, why populations fail to adapt and continue to expand their range, and what the long-term stability of range size is, are all fundamental questions in the ecology and evolution of ranges (Kirkpatrick & Barton 1997; Sexton et al. 2009; Gaston 2003; Louthan et al. 2015; Angert et al. 2020). Answering these questions has major implications for urgent areas of research, such as predicting species’ responses to climate change (Doak & William 2010) or the propensity of invasive species to rapidly expand their range (Alexander & Edwards 2010).

The simplest explanation for the persistence of a species’ range limits is that they reflect a species’ niche limits (Holt 2003; Hargraves et al. 2014); i.e., the species’ distribution is determined by the environment (Angert et al. 2020). When range and niche limits coincide, it suggests a failure of peripheral populations to adapt to local conditions and expand both niche and range. There are several competing evolutionary hypotheses that seek to explain the persistence of range edges due to a failure to locally adapt. Haldane (1956) and Mayr (1963) were among the first to suggest that asymmetric gene flow from central populations to the edge populations might limit the latter’s ability to adapt due to “gene swamping.” Gene swamping is expected to occur if migrants from central populations carry with them alleles adapted to “central” conditions but, due to antagonistic pleiotropy, maladaptive at the range edge and conditions beyond. If migration rates from the center populations are high enough, any adaptive alleles segregating in edge populations may become “swamped” by the influx of maladaptive alleles from migrants. Kirkpatrick & Barton (1997) formalized the gene swamping hypothesis by developing a quantitative genetic model across environmental clines of varying steepness, demonstrating that swamping could be a powerful mechanism maintaining species range edges. Gene swamping is a special case of the “migration load,” the reduction in fitness of a population due to the influx of migrant alleles maladapted to local conditions (Garcia-Ramos & Kirkpatrick 1997; Bolnick & Nosil 2007).

However, empirical evidence for the prevalence of gene swamping has been mixed. In a large review of the literature, Kottler et al. (2021) find little empirical evidence for gene swamping. Indeed, multiple studies indicate that increased gene flow to edge populations might have a positive effect (e.g., Sexton et al. 2011; Bontrager & Angert 2019). Range edges often have lower genetic diversity (Eckert et al. 2008; Pironon et al. 2016) and increased inbreeding depression (Arnaud-Haond et al. 2006; De Ryck et al. 2016) compared to core populations, both conditions that could be alleviated by increased migration. Therefore, Kottler et al. (2021) suggest that empirical evidence supports an alternative mechanism of range edge evolution: dispersal limitation (Sexton et al. 2009; Hargreaves et al. 2014). Under a dispersal limitation scenario, the lack of migrants to edge populations may be depriving them of the needed genetic variation to respond to strong selective pressures at the niche limit. As opposed to gene swamping, which predicts that increased migration from the center will hinder adaptation, the dispersal limitation hypothesis posits increased gene flow has the potential to rescue edge populations (Halbritter et al. 2015; Evans et al. 2014; Silva-Arias et al. 2017). However, due to the inherent complexity of species ranges, identifying range limits (e.g., Dallas et al. 2017; Pironon et al. 2017; Soberón et al. 2018) and the underlying processes generating those limits remains challenging (Kottler et al. 2021; Fristoe et al. 2023), especially in organisms not amenable to transplant experiments. Thus, an ideal system for investigating questions of range evolution would have a relatively simple range with well-defined edges.

In the ocean, particularly for pelagic organisms, an important cause of dispersal limitation may be the direction and strength of the dominant current regimes (Siegal et al. 2003). The average dispersal distance of a marine pelagic larva, for example, is a function of its settling time and current velocity (Pringle & Wares 2007; Wares & Pringle 2008). White et al. (2010) found that incorporating larval duration and directional oceanic currents dramatically improved model predictions of population differentiation in the Kellet’s whelk (*Kelletia kelletii*) over models relying on geographic distance alone.

Similarly, Dambach et al. (2016), investigating a deep-sea shrimp (*Nematocarcinus lanceopes*), identified patterns of relatedness between populations consistent with current connectivity in the Southern Ocean. Thus, marine ranges can be determined by complex abiotic and biotic factors that can cause mass movement of individuals in asymmetric ways. For example, in the North Atlantic, a northward current, the Gulf Stream, emerges from the Loop Current in the Gulf of Mexico, bringing warm water from the Caribbean, and travels up the coastline to Chesapeake Bay before diverting to the mid-Atlantic (**Fig. 1A**). A second regime, the Labrador Current, travels southward along Nova Scotia towards New England, bringing with it cold Arctic water, before also diverting eastward. Hence, along the North Atlantic coastline, two currents converge along the central latitude from opposite directions. For a marine organism distributed along the entire coastline and whose dispersal is largely dictated by the currents, this could generate a migration bias towards the center of the range. If the asymmetry in dispersal is strong enough, it might even suggest that range limits are not reflective of niche limits, but instead the lack of available migrants for successive colonization and further range expansion into otherwise habitable patches (Hargraves et al. 2014; Oldfather et al. 2019).

**Figure 1.**
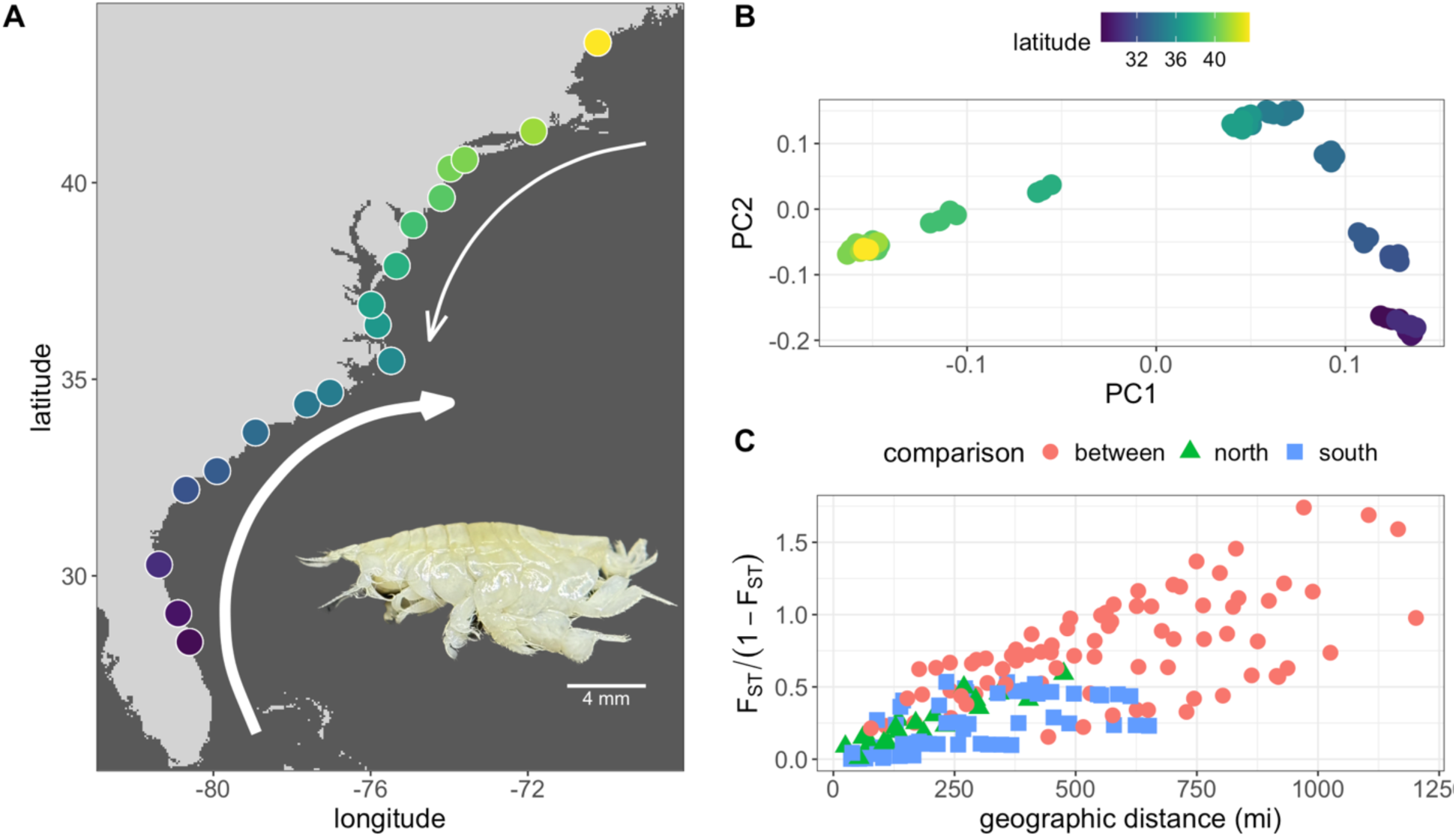
Sample locations and characterization of population genetic structure. A) Points are sampled localities, colored by latitude. Northbound arrows are the Gulf Stream, southbound are the Labrador Current, size differences represent different current intensities. Inset is a photograph of a shovel bug collected from New Jersey. B) PCA results; points are individuals, colored by the latitude of their sample site. C) Isolation-by-distance; points are sample localities plotted by their pairwise linearized *F*_ST_ against geographic distance in miles. Colors are comparisons within groups (north or south of the Chesapeake) or between them.

Small amphipod crustaceans in the family Haustoriidae, which inhabit the exposed sandy beaches of the western North Atlantic, are an excellent system in which to test questions about range evolution. Due to their life history (see *Methods*), haustoriid amphipods are expected to be relatively poor dispersers, and this has been largely confirmed in previous studies (Jones 2005; Hancock et al. 2019). In the western North Atlantic, *Haustorius canadensis* (Bousfield 1962) can be found on beaches from central Florida to southern Maine with some relict, isolated populations reported in the Gulf of Saint Lawrence (Bousfield 1962; 1965). Haustoriid amphipods have no prior common name; here we introduce the term “shovel bugs” (with reference to their modified shovel limbs) to describe haustoriids, and “Canadian shovel bug” to refer to *H. canadensis*. A curious feature of their southernmost distribution is the abundance of suitable sandy beaches farther south than they are found, with no Canadian shovel bug recorded south of Melbourne, Florida. The northern range edge, alternatively, might represent a hard environmental transition, as most of coastal Maine is composed of rocky outcrops with few fine sand beaches.

Given their relatively constrained niche (the intertidal of wave-exposed sandy beaches), the ease of identifying range boundaries, and the one-dimensional nature of their range, the Canadian shovel bug is an ideal organism for evaluating hypotheses of the evolutionary mechanisms determining range limits. If gene swamping is maintaining range edges in this species, we should detect a signature of asymmetric gene flow from the range center towards the edge. Furthermore, we expect edge populations to harbor the genetic signature of population sinks – lower genetic diversity relative to more central populations, and the genetic diversity they do possess should be a subset of that harbored in the center. Alternatively, range edges might be maintained by a failure to colonize habitable patches due to asymmetries in the dispersal medium. Because shovel bugs passively disperse via the currents, successively more distant patches in the opposite direction of the current flow may have fewer available migrants to colonize them, ultimately determining range limits. If dispersal limitation due to asymmetries in current regimes are driving the persistence of range edges, we expect to find patterns of asymmetric migration towards the range center. In addition, we would expect to identify population-clustering associated with the two major currents, and a breakdown of this clustering pattern at central latitudes where the currents converge.

In this study, we first characterize the population genetic structure of the Canadian shovel bug across its entire range in the North Atlantic. Next, we estimate genetic diversity and inbreeding across the range to test for patterns of reduced abundance along the range edges. Finally, we determine whether patterns of migration and colonization are consistent with either gene swamping or dispersal limitation as processes maintaining range edges. We find evidence for directional colonization from the periphery of the range toward the center, which we hypothesize is due to current regimes in the North Atlantic and consistent with the migration limitation hypothesis.

## 2 METHODS

### 2.1 Natural history of shovel bugs

Deemed “the most interesting group of amphipods,” shovel bugs are haustoriid amphipods that have highly derived morphological adaptations, including expanded articles on the 6th and 7th pereopods, extensive setal elaboration, and spiny, cup-like structures on their 3rd and 4th pereopods that they utilize as shovels (**Fig. 1A**) (Barnard 1969; Hancock & Wicksten 2018).

Shovel bugs are blind and entirely fossorial, living in the first 1–10 cm of sand and almost exclusively occupying the narrow swash zone (Croker & Hatfield 1980). They lack a pelagic larva; instead, females brood juveniles through the first few molts in their marsupium. Adult shovel bugs range in body size from 2–18 mm, sometimes within the same species, and past work has found strong correlations between body size and latitude (Hancock et al. 2021), though there may also be an effect of sand grain size (LeCroy et al. 2002; Hancock & Wicksten 2018). Several species may occupy the same beach, but there is evidence that they partition the beach by sex and by species to limit competition for space (Croker & Hatfield 1980).

The relationship of shovel bugs to the broader Amphipoda has been shrouded in mystery, with some hypothesizing a close kinship with other fossorial amphipods (Lowry & Myers 2017). However, recent molecular phylogenies have cast doubt on the taxonomic affinities between the fossorial amphipods, with multiple studies indicating shovel bugs are gammarids and that modifications to the fossorial lifestyle found across several families arose convergently (Verheye et al. 2016; Copilas-Ciocianu et al. 2020; Hancock et al. 2022).

### 2.2 Field collecting

Shovel bugs were collected from 18 sites from Cocoa Beach, Florida, USA to Crescent Beach, Maine, USA in May of 2023 (**Fig. 1A**; *Table S1*). Collections began in West Palm Beach, Florida, USA to identify the southernmost established population; traveling north from West Palm Beach, no specimens were located until Cocoa Beach, Florida. Similarly, sampling continued beyond Crescent Beach, Maine, but no samples were located further north (last attempted location was Popham Beach, ∼93 km north of Crescent Beach). Specimens were sifted from sand using a 435 *μ*m sieve plate, plucked from the plate with forceps, and preserved in 95% ethanol. Individuals were identified to species in the lab using a dissecting microscope and key from Hancock & Wicksten (2018), and then stored at -20°C prior to DNA extraction.

### 2.3 Molecular techniques

Whole genomic DNA was extracted from either the whole individual or from pereopods 4–7 using a Qiagen DNeasy Blood & Tissue Kit following manufacturer’s protocols. Library prep was performed using Restriction site Associated DNA sequencing (2RAD) following Bayona-Vásquez et al. (2019), with slight modifications. All 96 samples were quantified using a Qubit^TM^ 1X dsDNA High Sensitivity (HS) kit on a Qubit 2.0 fluorometer. Samples were then normalized to have 100ng per sample (5 µ*L*) on a 96 well plate, and then digested in a 15 µ*L* solution with 1.5 µ*L* 10X CutSmart Buffer (New England Biolabs, USA), with 10 units of EcoRI (0.5 µL), 10 units MseI (0.5 µ*L*), 3.5 µ*L* Invitrogen™ UltraPure™ DNase/RNase-Free Distilled Water, 2.5 µ*L* of EcoRI compatible adapter (2 µ*L*), and 0.25 µ*M* MseI compatible adapter (µ*L*) on a Eppendorf Mastercycler PRO S 6325 Thermal Cycler thermocycler at 37°C for one hour with the heated lid turned off. Immediately after digestion we added 5 µL of ligation mastermix composed of 2.75 µL of nuclease free water, 0.5 µL 10X ligation buffer (NEB), 1.5 µL 10 mM rATP (Promega), and 0.25 µL T4 DNA Ligase 400,000 U/ml to each sample. The ligation reaction was implemented on the same thermocycler as above for two cycles of 22°C for 20 min, 37°C for 10 min, followed by 80°C for 20 min to stop the enzyme activity. We then pulled 5 µL of each sample and split these into two microcenterfuge tubes and performed a 1.25X SpeedBead cleanup (Rohland and Reich, 2012).

To reduce PCR duplicates and add the iTru5–8N barcode (which has a segment of eight random base pairs) we implemented a single PCR cycle with six tubes each with a 50 µL reaction. Each reaction contained 10 µL nuclease free water, 25 µL KAPA HiFi hot start ready mix (Roche, USA), 5 µL iTru5–8N barcode (5 µM), and 10 µL of library and were added using a single cycle of 98°C for 1 min, 60°C for 30s, and 72°C for 6 min. All six PCR reactions were then pooled and a 1.5X SpeedBead cleanup was performed and resuspended in 30 µL of nuclease free water. Next, we split the library into three 55 µL reactions with three unique iTru7 adapters. Each reaction had 10 µL of nuclease free water, 25 µL KAPA HiFi hot start ready mix (Roche, USA), 5 µL P5 primer (5 µM) and 5 µL of each iTru7 (5 µM) added separately. The PCR was implemented on the thermocycler mentioned above under the following conditions: 95°C for 3 mins, 8 cycles of 98°C 20s, 60° 15s, 72°C 30s, and a final 72°C for 5 mins. All PCR reactions were pooled and then cleaned using a 1.5X SpeedBead cleanup and resuspended in 30 µL of nuclease free water.

We performed size selection on the pooled library using a Pippin Prep (Sage Science, USA) with internal standards on a 1.5% dye free cassette, target 350–480 bp range (mean= 415 bp). After size selection to enrich the library, we performed a PCR under the same conditions for the iTru7 reaction above, except with only four 50 µL reactions and 6 cycles, using both the P5 and P7 primers (5 µM). Finally, we performed a 1X SpeedBead clean up and quantified the library on the Qubit 2.0 to ensure that our library had a high enough concentration for sequencing. The library was then sequenced on 12.5% percent of a NovaSeq X lane at the University of Michigan Advanced Genomics Core.

Raw sequence reads were assembled and filtered using the de novo pipeline in STACKS v.2.67 (Catchen et al. 2013; Rochette et al. 2019). A custom *process_radtags* function was provided to us by J. Catchen to handle the unique FASTQ files of 2RAD, which include an additional adapter sequence. Reads were assembled using the de novo pipeline (Catchen et al. 2011); briefly, stacks were built in *ustacks* with a maximum nucleotide distance allowed of 2 and minimum number of reads needed to seed a stack of 3. Stacks were merged allowing a maximum of 2 gaps and 6 mismatches. Next, we used the *populations* function, filtering out alleles that did not occur in at least 10 populations and keeping those that were present in 75% of individuals per population. Lastly, we randomly selected one SNP per RAD locus (--write-random-snp). Sequences were output as two VCF files, one that included only variant sites and a second that contained both variant and monomorphic sites. VCF files were additionally filtered using *vcftools* v.1.16 (Danecek et al. 2011); first, individuals with >80% missing data were removed. Next, we filtered out loci with more than two alleles (--max-alleles 2) and with a minor allele frequency of less than 0.05 (--maf 0.05). We then filtered out loci with a minimum mean depth of less than 20 (--min-meanDP 20) and removed all indels (--remove-indels). After filtering, we evaluated the relationship between mean depth, estimated heterozygosity, and distance from the range center via linear regression to determine if mean depth might be a confounding factor.

### 2.4 Population genetic analysis

To evaluate potential patterns in genetic diversity, we estimated individual-level heterozygosity using *pixy* (Korunes & Samuk 2021) on the VCF that included invariant sites. The inbreeding coefficient (*F*; Wright, 1931), estimated as the difference between the expected and observed homozygosity per individual, was estimated using *vcftools* (- -het).

To test for geographic structure, we first used *pixy* to estimate population-level pairwise *F_ST_* (Weir & Cockerham 1984), which we then linearized and regressed against geographic distance to test for isolation-by-distance (Wright 1943; Rousset 1997). Next, we performed a principal component analysis (PCA) to identify signs of genetic clustering using *VCF2PCACluster* (He et al. 2024). We then performed two cluster analyses. First, we used *fastSTRUCTURE* (Raj et al. 2014) with *k* (the number of discrete, ancestral clusters) from 2–5. Next, to account for continuous spatial differentiation, we ran *conStruct* (Bradburd et al. 2018), varying *k* from 1–4. For each value of *k*, we performed 5 independent MCMC chains each with 2000 iterations. We then reported results from the chain with the highest mean posterior probability. As an additional test of whether identified clusters and admixed individuals were the result of increasing gene flow towards the range center or continuous isolation-by-distance, we also performed a triangle plot analysis using *triangulaR* (Wiens & Colella 2024a). In the presence of directional gene flow, individuals with a hybrid index approaching 0.5 should have concomitantly high interclass heterozygosity. However, a continuous population characterized solely by isolation-by-distance will display a high hybrid index but no increase in interclass heterozygosity (Wiens & Colella 2024b).

We evaluated potential barriers to migration using the program *FEEMS* (Marcus et al. 2021), which estimates an effective migration surface using edge-specific parameter estimation in a pseudo-likelihood framework. Lastly, we tested for a signature of range expansion along the periphery of the population using Peter & Slatkin’s (2013) directionality index, *ψ*, as implemented in *rangeExpansion* (https://github.com/BenjaminPeter/rangeexpansion). The directionality index is calculated by comparing the difference in allele frequencies between pairs of populations, conditioned on the allele being present in both populations. Large source populations typically contain many rare alleles. When a new population is founded from this larger source population, these same alleles, if they have not been lost due to drift, are likely to drift to higher frequencies in the new population. Thus, successively more recently founded populations in an expanding range will have increasingly positive values of *ψ*, permitting the identification of both the origin and the direction of the range expansion (Peter & Slatkin 2013; 2015). Following Peter & Slatkin (2015), the directionality of range expansion is inferred by summing across the columns of the matrix of pairwise estimates of *ψ* for each population (by row), where the value of *ψ* is the strength of the signal and the sign is the direction. In populations at migration-drift equilibrium in continuous space, simulations show that *ψ* should be zero (Peter & Slatkin 2013). Therefore, if *ψ* is zero, there is no marked departure from migration-drift equilibrium and hence no signature of reduced fitness or recurrent colonization in edge populations. However, if gene swamping is driving edge dynamics, we expect *ψ* should be positive in the direction of the edge. Alternatively, if dispersal limitation, driven by directional currents, is the primary cause of range limits, then *ψ* should be negative at the edges and increasingly positive toward the range center.

### 2.5 Simulations

Because the expansion statistic *ψ* is influenced by relative rates of drift, the inferred direction of population expansion, as given by *ψ*, may be influenced by spatial clines in population density even under scenarios with isotropic dispersal. To determine whether the results of our *ψ* analyses are robust to spatial variation in population density (see Results/Discussion), we conducted forward-time simulations using SLiM v.4.0.1 (Haller & Messer 2023). We established a 10-deme stepping-stone model with four different range scenarios: 1) equal population size per deme with symmetrical migration; 2) equal population size per deme with asymmetrical migration from the edge populations towards the center; 3) an abundant center model in which the central demes had a fivefold higher census population size than the edge, with symmetric migration; and 4) abundant center with asymmetric migration from the edges inward. For the asymmetric migration scenarios, migration rates from the edge towards the center were twice that of center to edge, and for scenarios with an abundant population center, population size decayed linearly towards the edge. In addition, we performed simulations with increasing differences in population sizes between the central and edge populations, with equal migration, to evaluate how these relative differences in genetic drift might drive patterns in *ψ*. We simulated ratios of population size between the center and edge of 5:1, 25:1, and 50:1 and performed 10 replicates for each scenario. For each ratio, population size decreased linearly, but more precipitously at higher ratios to balance maintaining constant deme number and computational efficiency. Individuals were diploid and hermaphroditic, and we simulated a 10 Mb chromosome with a recombination rate of 1×10^−8^ per base pair. Simulations were run for 100,000 ticks (>10*N*) to reach demographic equilibrium, and tree sequences were recorded for subsequent processing using *pyslim* v.1.0.1 (Kelleher et al. 2018) and *tskit* v.0.5.3 (Kelleher et al. 2018). For tree sequences with more than one root, we used recapitation to simulate a neutral coalescent history until only a single root remained; mutations were added at a rate of 1×10^−7^ per base-pair to the trees using *msprime* v.1.2.0 (Kelleher et al. 2016). Next, we randomly sampled five individuals per deme and output their sequences in VCF format. From here, we trimmed the VCFs to only include biallelic SNPs, and converted them to BED files using *plink* v.1.9 (Purcell et al. 2007). To use the *rangeExpansion* package, we converted the individual coordinates from SLiM, which were in discrete demes, into Earth coordinates as required by *rangeExpansion*; thus, we chose 10 coordinates 5° apart in latitude. Because the units of distance are arbitrary in SLiM, we needed only to ensure that deme 1 is closest to deme 2, etc., to adequately capture the SLiM environment in an Earth coordinate system. We then calculated *ψ* for each simulation to see how the different range scenarios might impact inferences made from *ψ*.

## 3 RESULTS

### 3.1 ddRAD assembly

A total of 2,605,149 loci passed filtering in STACKS; the mean effective per-sample coverage was 23.0 with a standard deviation of 13.6, a minimum of 4.0, and a maximum of 67.4. The mean number of sites (bp) per locus was 220.1. After filtering for biallelic SNPs and missingness in *vcftools*, the final dataset included 7,405 SNPs. We found a significant positive relationship between mean depth per individual and our estimates of heterozygosity from *pixy* (*P* < 5.3e–7, R^2^ = 0.27, *β* = 8.77e–6; *Table S2*); in addition, we found a significant negative relationship between mean depth and the distance from the center of the range (taken as 36° latitude; *P* < 0.002, R^2^ = 0.10, *β* = –0.051) (see *Table S2*). To account for the potential confounding between measures of heterozygosity, mean depth, and their relationship with the distance from the range center, all subsequent multivariate linear regressions were performed with mean depth as an additional predictor. To determine if a few individuals with high mean read depth were driving the observed patterns, we re-tested for a significant correlation between depth and heterozygosity after dropping all individuals with a mean depth greater than 30. While a significant correlation between heterozygosity and mean depth remained (*P* < 2.7e–5; R^2^ = 0.38, *β* = 3.1e–5), it was no longer significantly correlated with distance from the center (*P* = 0.57; R^2^ = –0.02, *β* = –2.03) (*Fig. S1*).

### 3.2 Genetic structure

The PCA revealed relatively continuous structure, with a possible break between populations north and south of the Chesapeake (**Fig. 1B**), but there were no obvious discrete breaks identified in patterns of isolation-by-distance when regressing linearized *F_ST_* against geographic distance (**Fig. 1C**). Results of the *fastSTRUCTURE* analysis supported three geographically restricted population clusters (*k* = 3) as the model that maximized the marginal likelihood – one from Florida to South Carolina, a second from North Carolina to Virginia, and a third that included the remaining sample sites from northern Virginia to Maine (*Fig. S2*). However, *conStruct*, which allows for continuous structure to be accommodated within discrete populations, supported *k* = 2. Identified clusters corresponded to the dominant current regimes in the Atlantic with increasing admixture towards the center of the range (**Fig. 2**). This result was echoed by the triangle plot analysis of the *conStruct* results, which found increasingly elevated interclass heterozygosity at central latitudes (*Fig. S3*).

**Figure 2.**
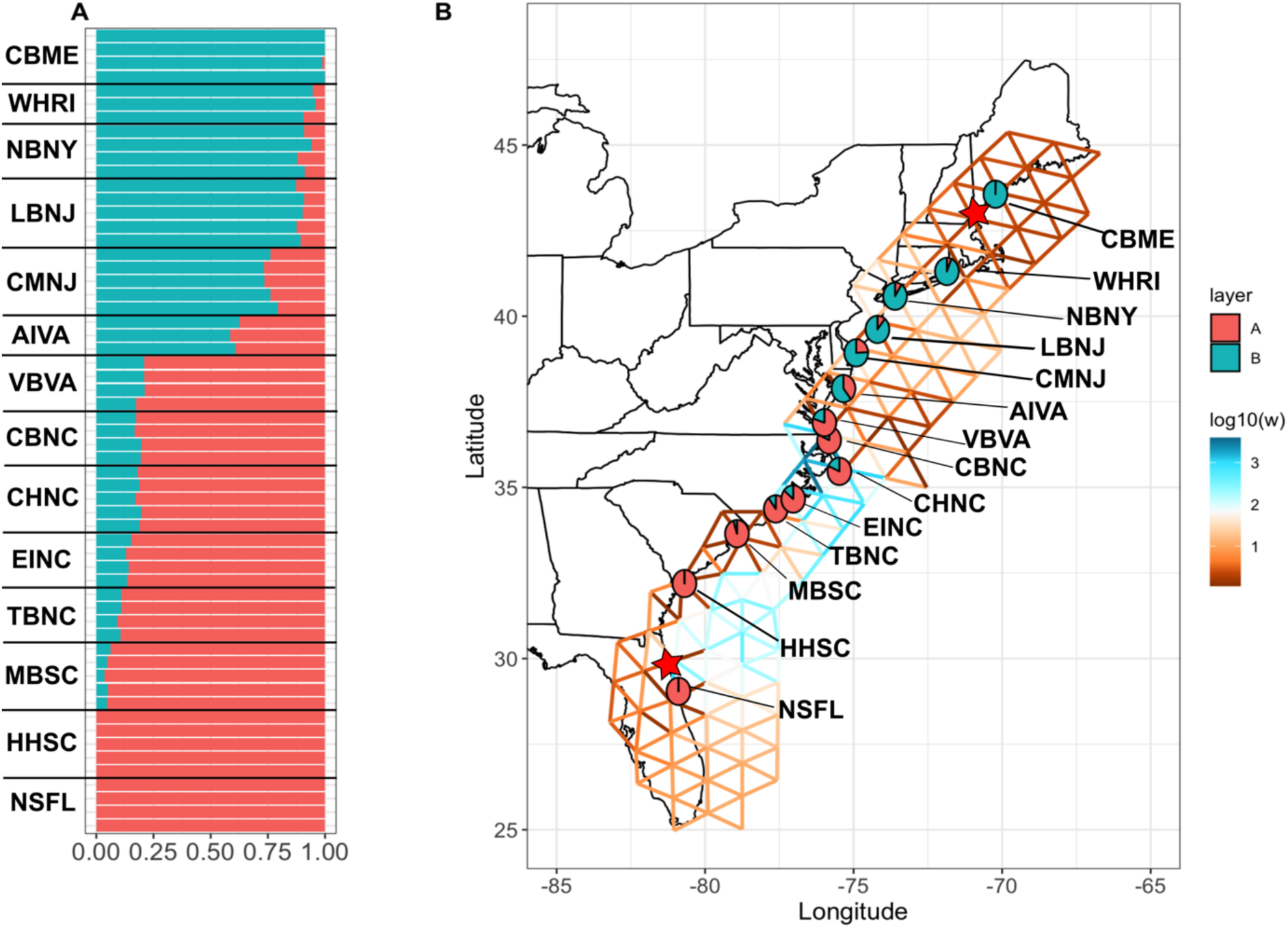
Admixture and effective migration surface. A) Clustering (*conStruct*) analysis results using *k* = 2. B) Effective migration surface from *FEEMS*. Cooler colors represent regions of higher migration, warmer indicate potential barriers. Pies represent the admixture proportions inferred from *conStruct* averaged across all individuals sampled at a locality. Stars represent inferred origins of the population expansion from the *ψ* analysis. Population abbreviations are defined in *Table S1*.

### 3.3 Patterns of diversity

After incorporating sequencing depth as a possible confounding effect, measures of individual heterozygosity were significantly predicted by the distance from the center of the range (*P* < 3.79e–16, *β* = –1.5e–6; mean depth *P* = 5.1e–5, *β* = 5.38e–6). This relationship remained significant after dropping individuals with mean depth greater than 30 (*P* < 0.0009; R^2^ = 0.25, *β* = –1.2e–6) (*Fig. S4*). Similarly, measures of inbreeding were the highest at the range edges, declining as a function of distance from the center (**Fig. 3B**). These patterns are consistent with field observations: the ease of collecting and observed highest abundance was in the central range locations and shovel bugs at the edge sample locations were rarer and thus more difficult to collect. However, we note that while distance is a much stronger predictor of patterns of genetic diversity and inbreeding, mean sequencing depth was still a significant predictor in each model (*Table S2*).

**Figure 3.**
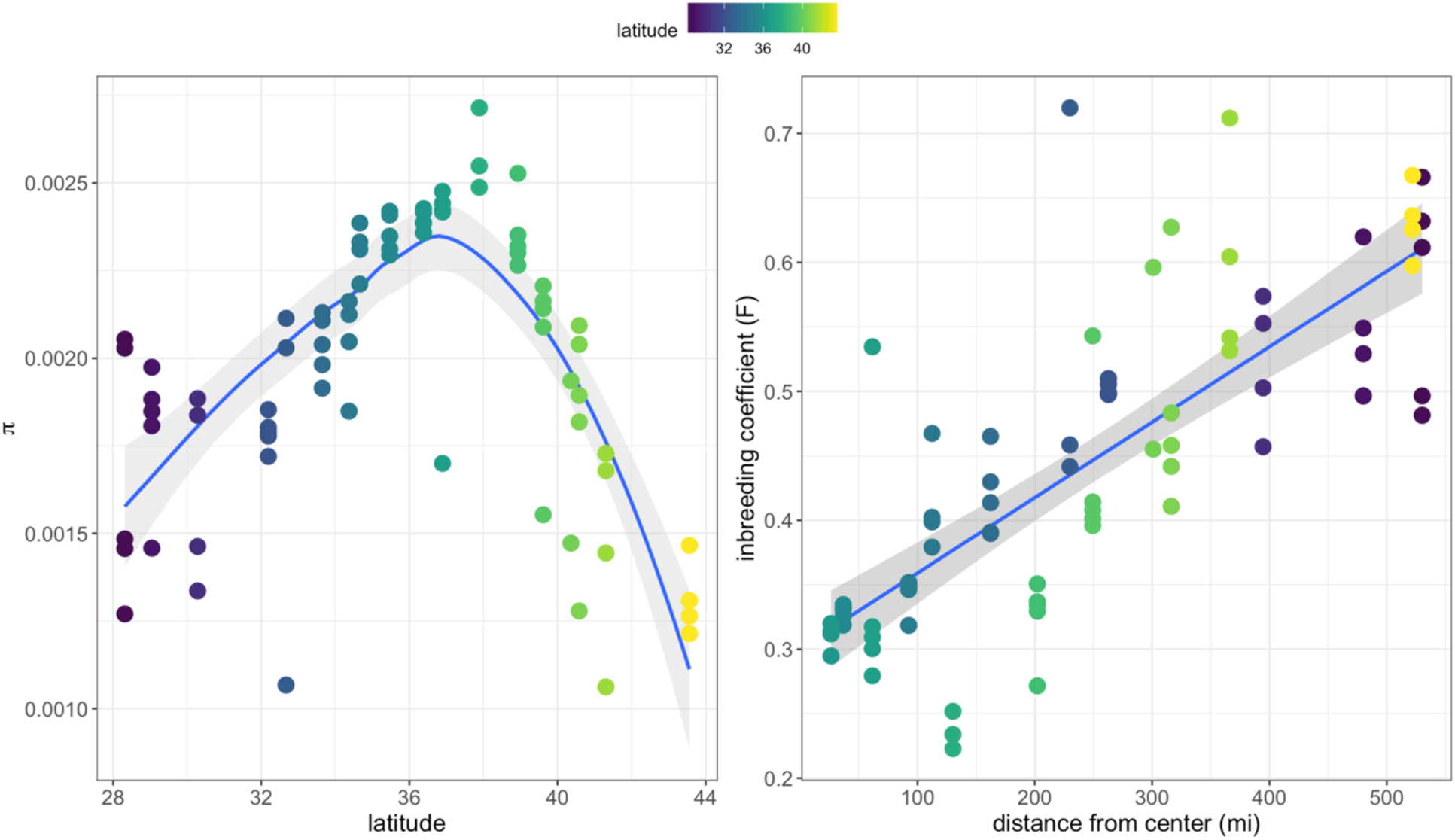
Patterns of genetic diversity. A) Individual genetic diversity (π) as a function of latitude. B) Inbreeding coefficient as a function of the distance from the center of the range, taken as 36° latitude, in miles. Points are colored by the latitude of their sample location as in Fig. 1A.

The effective migration surface estimated from *FEEMS* suggested elevated levels of dispersal from northern Florida to Virginia along the path of the Gulf Stream and reduced effective migration towards the edges (**Fig. 2B**). In accordance with results from the PCA, the migration surface indicated a strong depression in effective migration across the Chesapeake, which suggests it might act as a partial barrier to gene flow.

### 3.4 Directionality index & simulations

The inferred origin of the range expansion of the entire sample of individuals, as indicated by the lowest value of *ψ*, was 29.9°N in northern Florida (**Fig. 4A**). When the data were subset into two regions, South and North (based on the *conStruct* analysis), the inferred origins were at 29.9° and 43.6° latitude, respectively. The sum of *ψ* for each population resulted in a pattern of increasing *ψ* towards the range center, mirroring the increase in heterozygosity (**Fig. 4A**). This pattern of *ψ* superficially suggests a range expansion from the edges toward the center of the range that follows the dominant current regimes of the Gulf Stream running south to north and the Labrador Current running north to south, both converging in the center of the range (**Fig. 1A, 4A**). This pattern of expansion is in opposition to expectations from the gene swamping hypothesis and suggests that asymmetric migration due to the currents might be maintaining range edges, consistent with the migration limitation hypothesis.

**Figure 4.**
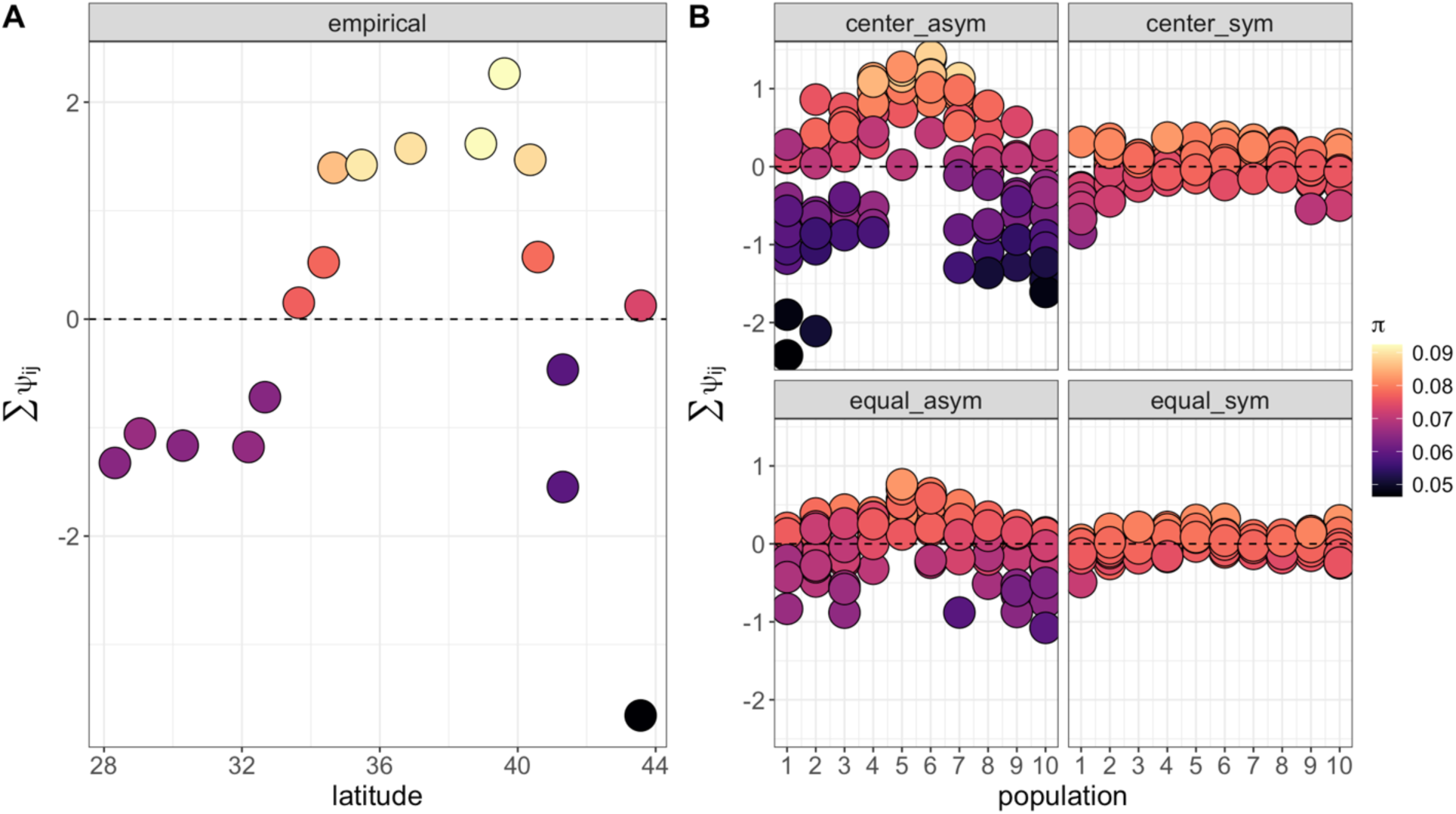
Directionality index (*ψ*) results. A) Empirical estimates of *ψ*, points are the summed values of *ψ* for all comparisons of population *i* to populations *j*. B) Simulation results for each stepping-stone model: “center_asym” is abundant center with asymmetric migration; “center_sym” is abundant center with symmetric migration; “equal_asym” is equal deme size with asymmetric migration; and “equal_sym” is equal deme size with symmetric migration. Each point per population is a different replicate. Points are colored by estimated genetic diversity (π) of the sampled locality. Dashed line is when *ψ* = 0, which is expected under equilibrium isolation-by-distance.

Simulation results from SLiM supported this interpretation. Across replicates, the strongest divergences from *ψ* = 0, which is expected at migration-drift equilibrium, occurred in simulations with asymmetric migration from the edges toward the center, with the most striking inflation in the scenario in which there is both an abundant center and asymmetric migration (**Fig. 4B**). While there was some deviation from zero along the edges in scenarios with symmetric migration in both the equal and center-abundant scenarios, they did not systematically produce a positive *ψ* in the center as in the case of asymmetric migration (**Fig. 4B**). Furthermore, increasing the population size discrepancy between the center and the edge did not consistently lead to an increase in central estimates of *ψ* (*Fig. S5*), indicating that asymmetric migration, not increased drift along the edges, is the dominant force shaping the observed dynamics.

## 4 DISCUSSION

The persistence of range edges despite the potential for local adaptation to marginal conditions and resulting continued expansion is an enduring evolutionary mystery. Several hypotheses have been proposed to explain the finite nature of species ranges, principle among these gene swamping (Mayr 1963; Kirkpatrick & Barton 1996) and dispersal limitation (Sexton et al. 2009; Hargraves et al. 2014). In this study, we investigated the suitability of these hypotheses to explain range evolution of the Canadian shovel bug (*Haustorius canadensis*; Bousfield 1962), which is distributed along the Eastern Seaboard of North America.

Past work suggests that shovel bugs have the potential for large amounts of cryptic diversity and are sensitive to environmental barriers (Jones 2005; Hancock et al. 2019). Thus, our first aim was to characterize the population genetic structure of the Canadian shovel bug. While we found that patterns of linearized *F_ST_* were consistent with a single continuous population, the PCA and *conStruct* results suggested the presence of two populations associated with each current regime. In addition, we found evidence for resistance to migration across the Chesapeake Bay. Shovel bugs of the genus *Haustorius* specialize on open, wave-exposed beaches in which the sand is highly oxygenated (Bousfield 1970), and rarely occur in low salinity, brackish bays. While the Chesapeake Bay was formed by an impact crater during the Eocene (Ivany et al. 2000), and this event has been postulated to have contributed to haustoriid diversification in North America (Hancock et al. 2022), its recent form as a large bay is more recent (around the end of the Last Glacial Maximum ∼10,000 years ago (Colman & Mixon 1988)). Thus, the bay has the potential to be acting as a partial barrier to dispersal and gene flow. However, because the isolation-by-distance regression did not support a discrete break (**Fig. 1C**; *Fig. S3*), the barrier may be weak or has occurred recently enough that it has not yet influenced genome-wide patterns of allele frequency divergence, which can lag behind other population summary statistics (Barton et al. 2013; Ringbauer et al. 2018).

Once the genetic structure of shovel bugs was characterized, our next aim was to determine if range edges have reduced genetic diversity and increased inbreeding relative to central populations. Past theoretical (Polechová & Barton 2015) and empirical (Eckert et al. 2008; Pironon et al. 2016) work suggest that genetic diversity generally decreases as a function of distance from the range center when the environment changes gradually across space. Consistent with this past work, we found genetic diversity and inbreeding strongly correlated with distance from the range center (**Fig. 3**); edge populations harbor the lowest genetic diversity and highest rates of inbreeding. This pattern is broadly consistent with the “abundant center hypothesis” (Grinnell 1922; Hengeveld & Haeck 1982; Brown 1984), which predicts that population abundance is highest in the core of the range and decreases monotonically towards the edge. However, the generality of the abundant center hypothesis remains hotly contested, as there is little empirical evidence that population abundance consistently declines from the core to the edge (Dallas et al. 2020; Santini et al. 2019; Sagarin et al. 2006). While we observed higher densities of shovel bugs in the central sample sites, anecdotally consistent with the abundant center hypothesis, future work would benefit from quantitative estimates of census abundance across the Canadian shovel bug’s range.

Reduced genetic diversity and elevated rates of inbreeding in peripheral populations might indicate a reduced adaptive capacity (Angert et al. 2020; Kottler et al. 2023). During a range expansion, this can be driven by allele “surfing,” which describes the process by which genetic drift is increased along an expanding front due to reduced population densities, allowing alleles to reach high frequencies (Excoffier & Ray 2008). When a range size is stable, reduced adaptive capacity might result from gene swamping or dispersal limitation (Angert et al. 2020); both are expected to reduce population density on the range periphery. If range limits are being determined by gene swamping, we expected to infer asymmetric migration from the center toward the edge, with edge populations harboring a subset of the genetic diversity of central populations. Furthermore, we predicted that the shared alleles with central populations would be at higher frequency on the edge, consistent with repeated founder events at increasingly peripheral populations. Conversely, if range limits are being determined by dispersal limitation, we expected to infer a net direction of movement towards the range center from the edges, which might be reducing dispersal to peripheral populations. This is due to the dominant current regimes in the North Atlantic flowing in opposite directions, converging at a central latitude in the Chesapeake Bay region (**Fig. 1A**). In addition, if dispersal limitation is driving the persistence of range edges, we expected increasing admixture proportions towards the center of the range where the Gulf Stream and Labrador Current converge.

Our results strongly support the finding that dispersal limitation is the dominant mechanism shaping the persistence of range edges in shovel bugs. Our spatial clustering analysis indicated there are two populations of the Canadian shovel bug associated with each of the current regimes in the North Atlantic, with admixture proportions increasing toward the range center (**Fig. 2**). We also identified reduced effective migration on the range periphery (**Fig. 2B**). Interestingly, we observed that effective migration is generally higher south of the Chesapeake than north, which is consistent with the greater velocity of the Gulf Stream relative to the much weaker Labrador Current (**Fig. 1**). Furthermore, we detected a linear increase in *ψ* towards the range center from each edge. The standard interpretation of this index would imply a range expansion from the periphery towards the center. However, this is contradictory to patterns of inbreeding and genetic diversity, which clearly indicated that drift is likely more dominant on the range edge (**Fig. 3**). A possible explanation for these conflicting patterns is that the *ψ* index is influenced by asymmetric dispersal, and, if so, the observed directionality would support the dispersal limitation hypothesis.

To evaluate whether inferences made on the basis of the *ψ* index could be affected by heterogeneity in density as well as migration, we conducted forward-time, individual-based simulations intended to tease apart the impacts of differences in population size and migration (**Fig. 3B**; *Fig. 5*). Consistent with our explanation above, we found that *ψ* was highly sensitive to asymmetric migration and would infer a population expansion in the direction of the range center even in equilibrium (i.e., non-expansion) scenarios. In addition, this bias was strongest when under the combined influence of asymmetric migration and abundant center dynamics (**Fig. 3B**), but, importantly, did not occur in the presence of differences in population size alone, at least not at the ratios we tested (*Fig. S5*). This suggests that asymmetric migration may be playing the dominant role in shaping patterns in the SFS that are being identified by the *ψ* index.

The *ψ* index is a popular statistic for testing for range expansions and has been applied to a diverse array of different organisms (Manuel et al. 2016; Zhan et al. 2014; Fifer et al. 2022). In this study, we uncovered a potential confounding influence on *ψ* that is likely to impact any organism with high asymmetric migration. While asymmetric gene flow has been identified in terrestrial organisms (Fedorka et al. 2012; Oswald et al. 2017), the taxa most likely to display directional dispersal are aquatic organisms with passive dispersal via river flow or oceanic currents (Siegal et al. 2003; Pringle & Wares 2007; Morrissey & de Kerckhove 2009). Past work has shown that directional dispersal in the ocean can mislead many population genetic analyses, including the inference of random dispersal via patterns of isolation-by-distance and the probability of coalescence (Nagylaki 1977; Lenormand 2002; Paz-Vinas et al. 2013). Using a simulation approach, we uncovered that asymmetric migration can also bias the detection of a range expansion, potentially generating a false positive signature when, in fact, the population is in demographic equilibrium.

In addition to the impact asymmetric migration has on the inference of a range expansion, certain biased dispersal patterns may themselves be capable of generating an abundant center pattern. Specifically, asymmetric migration from the edges towards the range center can maintain high abundance in the center, causing it to act like a population sink, while colonization towards increasingly distant patches becomes less likely as the number of available migrants dispersing away from the center becomes increasingly small. Our findings in this system suggest that this may be an underappreciated form of migration limitation (Sexton et al. 2009; Hargraves et al. 2013), in which the rate of extinction in peripheral patches exceeds that of colonization; the cause of this decreased migration is asymmetries in the probability of migration due to the dispersal medium itself.

The evolutionary processes that maintain range edges, especially in systems in which the environmental change is clinal, are still subjects of active debate (Kottler et al. 2023). We uncovered strong support for the hypothesis that range edges in this system are maintained by dispersal limitation, itself driven by a combination of passive dispersal and directional currents reducing the number of available migrants to colonize peripheral patches. We suggest this mechanism for generating an apparent abundant center might be widespread in coastal marine taxa, especially those with passive dispersal, as oceanic currents regularly converge within species ranges. Furthermore, focusing on asymmetric migration via directional water or wind currents may help clarify what maintains range edges in cases where the fundamental niche appears to exceed the range limit.

## DATA AVAILABILITY

Bioinformatic and analysis pipelines, including filtered VCF data objects, are available at https://github.com/zachbhancock/asym_gene_flow_proj. Raw sequences were deposited to GENBANK (ascension PRJNA1183394).

## AUTHOR CONTRIBUTIONS

ZBH and GSB conceptualized and designed the study. ZBH carried out the field collections. ZBH and PLW performed the laboratory work. ZBH and NEA carried out the bioinformatics and analyzed the data. All authors wrote and edited the manuscript.

## ACKNOWLEDGEMENTS

Thanks to members of the Bradburd lab who provided helpful comments on this manuscript. Thanks also to Kristen Wacker, Teresa Pegan, and Benjamin Peter for thoughtful discussion and feedback. We are indebted to Leonard Jones who coined the common name “shovel bug” for haustoriid amphipods, and proudly use it in print for the first time here. May it catch on. Lastly, special thanks to Daniel Stern Cardinale and Robert Joel Duff for their hospitality and conversations while ZBH was collecting. Research reported was supported by the National Institute of General Medical Sciences of the National Institutes of Health (NIH) under award R35GM137919 (awarded to GSB). The content is solely the responsibility of the authors and does not necessarily represent the official views of the NIH.

