## Supplementary Material for "Asymmetric gene flow maintains range edges in a marine invertebrate"

**SUPPLEMENTARY MATERIALS**

**Tables**

*Table S1.* Sample locations.

| **Population** | **Label** | **Latitude** | **Longitude** | **# of samples** |
| --- | --- | --- | --- | --- |
| Cocoa Beach, FL | CBFL | 28.316 | -80.607 | 5 |
| New Smyrna, FL | NSFL | 29.043 | -80.897 | 5 |
| Jacksonville Beach, FL | JBFL | 30.282 | -81.386 | 4 |
| Hilton Head, SC | HHSC | 32.192 | -80.695 | 5 |
| Folly Beach, SC | FBSC | 32.668 | -79.907 | 3 |
| Myrtle Beach, SC | MBSC | 33.649 | -78.927 | 5 |
| Topsail Beach, NC | TBNC | 34.371 | -77.62 | 4 |
| Emerald Island, NC | EINC | 34.66 | -77.034 | 4 |
| Cape Hatteras, NC | CHNC | 35.469 | -75.48 | 5 |
| Currituck Beach, NC | CBNC | 36.381 | -75.824 | 4 |
| Virginia Beach, VA | VBVA | 36.892 | -75.985 | 5 |
| Assateague Island, VA | AIVA | 37.888 | -75.341 | 3 |
| Cape May, NJ | CMNJ | 38.93 | -74.912 | 5 |
| Long Beach, NJ | LBNJ | 39.615 | -74.197 | 5 |
| Sea Bright, NJ | SBNJ | 40.362 | -73.972 | 2 |
| Nickerson Beach, NY | NBNY | 40.585 | -73.605 | 5 |
| Watch Hill, RI | WHRI | 41.308 | -71.859 | 4 |
| Crescent Beach, ME | CBME | 43.564 | -70.224 | 4 |

*Table S2*. Regression models.

| **Model** | P-value | R^2^ | Mean_depth *p*-value | distance *p*-value |
| --- | --- | --- | --- | --- |
| heterozygosity~mean_depth | **5.281e–7** | 0.277 |  |  |
| mean_depth~latitude | 0.4968 | –0.007 |  |  |
| mean_depth~distance_from_center | **0.0026** | 0.1026 |  |  |
| heterozygosity~distance_from_center + mean_depth | **3.794e–16** | 0.6066 | **5.14e–5** | **1.35e–11** |
| F~distance_from_center + mean_depth | **< 2.2e–16** | 0.6661 | **0.00019** | **6.98e–15** |
| F_ST_~geographic_distance | **< 2.2e–16** | 0.5698 |  |  |

*Figure S1.* Correlations between mean depth, π, and the distance from the center of the range after dropping all individuals with mean depth > 30. A) Mean depth and π; *P* < 2.78e–5. B) Distance from the center in miles against mean depth; *P* = 0.575.

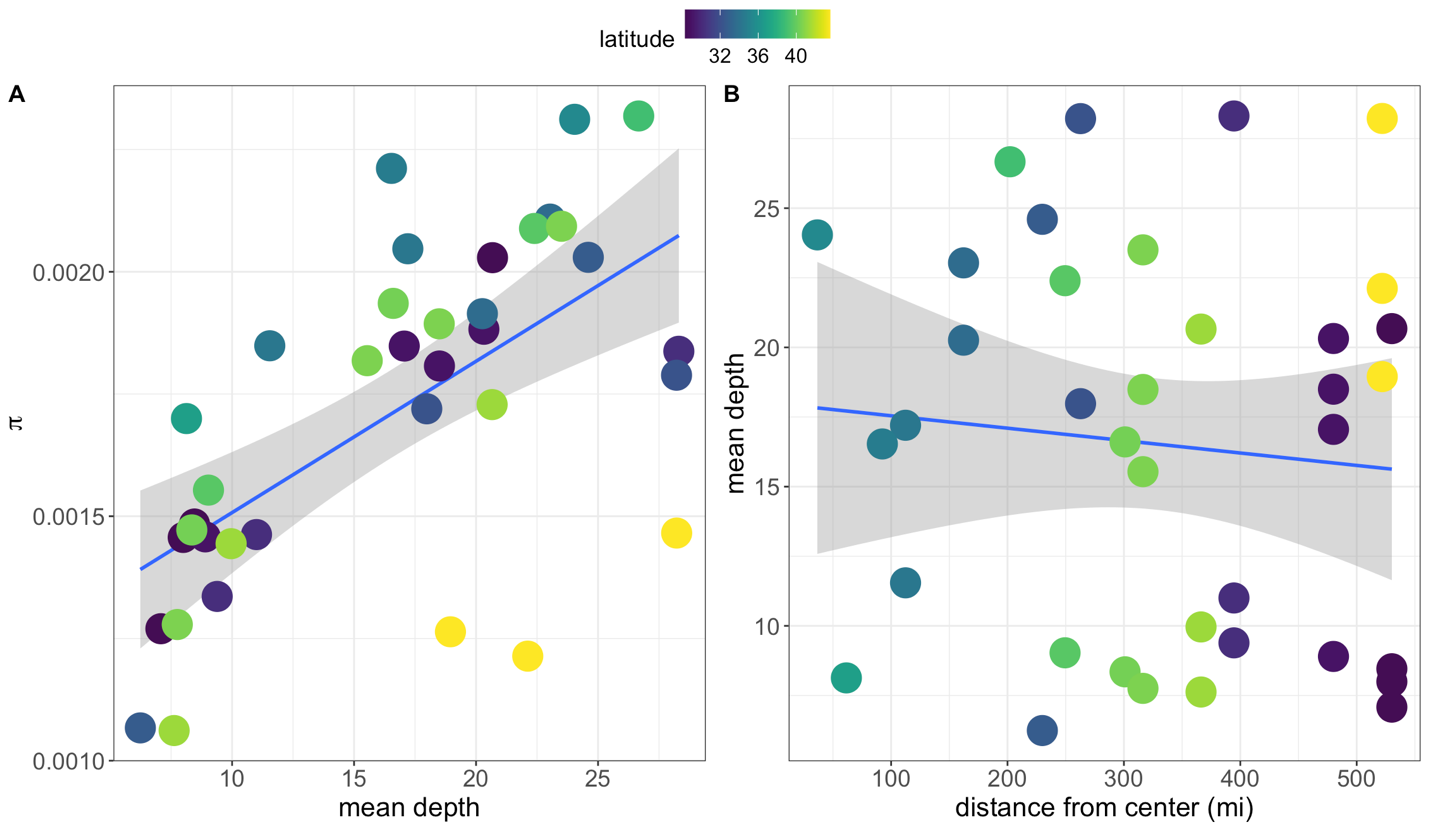

*Figure S2. fastSTRUCTURE* results for K=2–3. Each bar is an individual, samples are ordered by latitude.

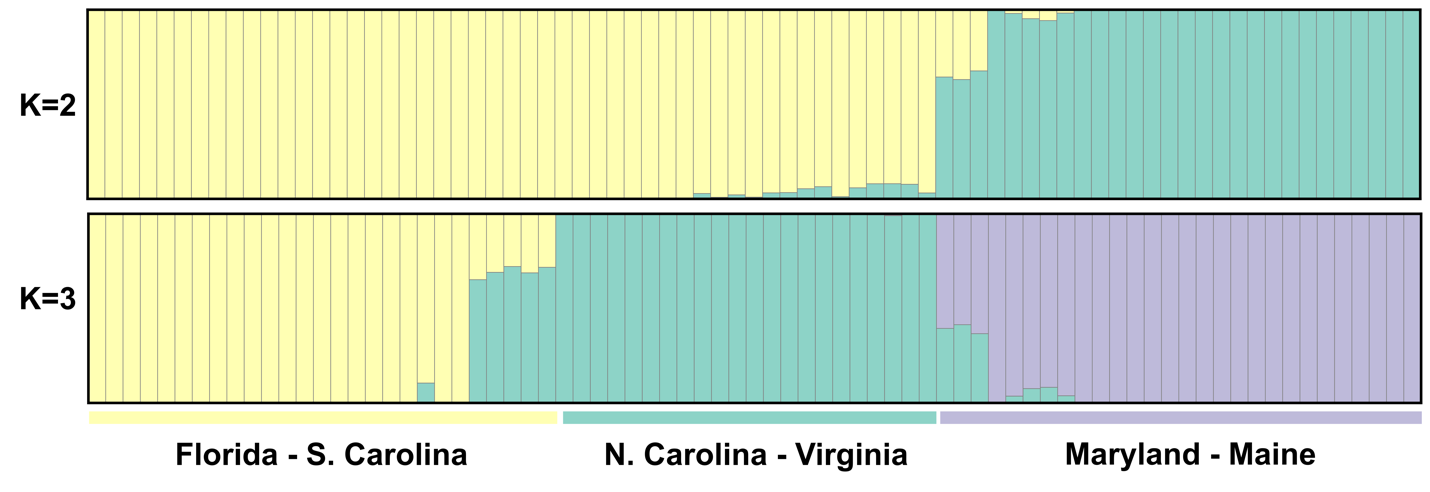

*Figure S3*. Triangle plot. Each point is an individual colored by sample location (“pop”). Notice that despite the increase in interclass heterozygosity towards the range interior, all population still fall well below the expected admixture proportions if there was hybridization in the presence of discrete structure (the dotted line).

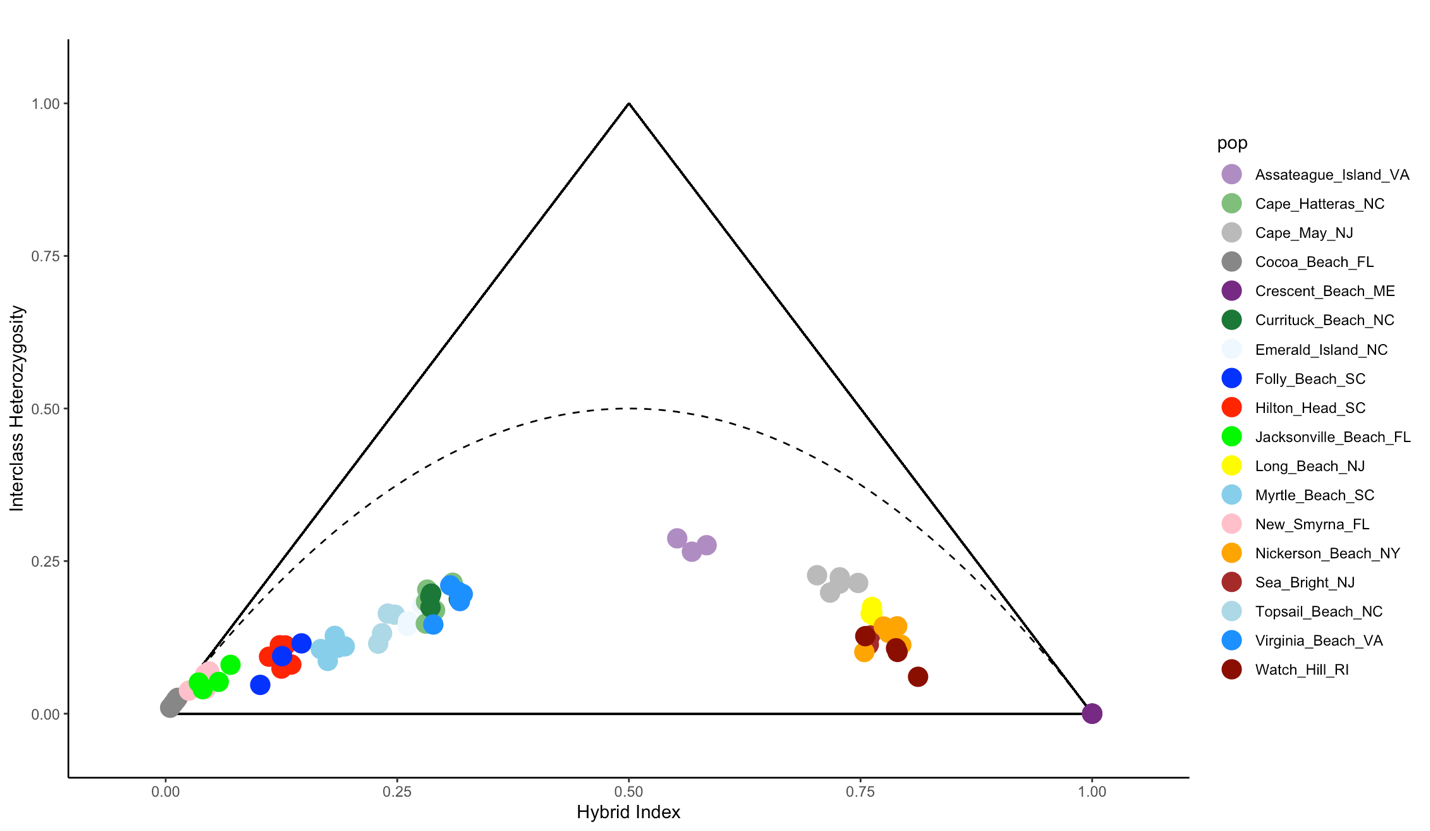

*Figure S4*. Relationship between π and the distance from the range center (in kilometers) after dropping all individuals with mean read depths greater than 30 (*P* < 0.0009; R^2^ = 0.25).

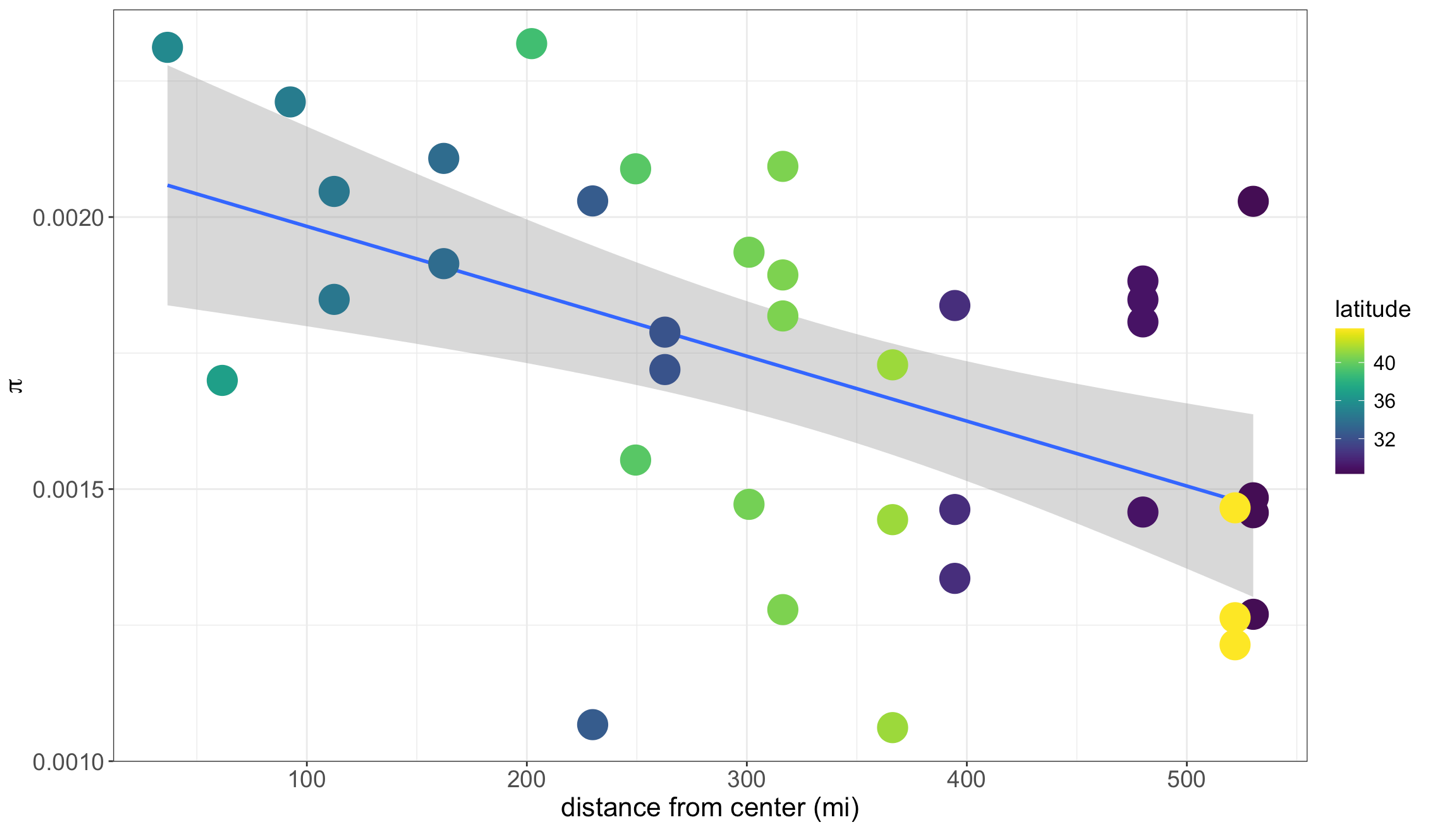

*Figure S5*. Simulation results for the directionality index ($\psi$) across three size disparity scenarios for the abundant center with symmetric migration model. Each point per population represents a different replicate; points are colored by estimated heterozygosity (π).

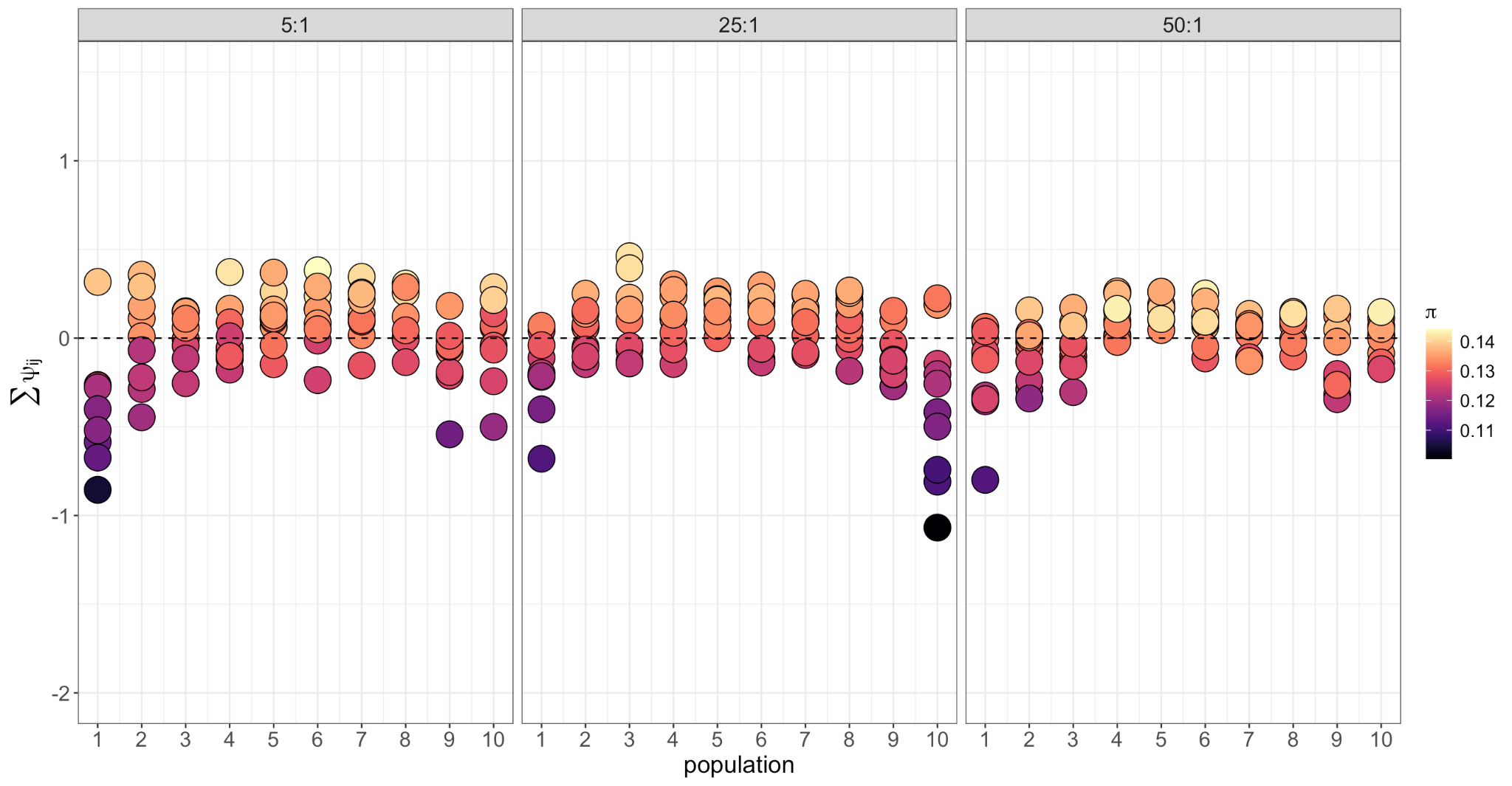
